# Quantifying and Visualizing Uncertainty for Source Localization in Electrocardiographic Imaging

**DOI:** 10.1101/2022.09.02.506414

**Authors:** Dennis K. Njeru, Tushar M. Athawale, Jessie J. France, Chris R. Johnson

## Abstract

Electrocardiographic imaging (ECGI) presents a clinical opportunity to noninvasively understand the sources of arrhythmias for individual patients. To help increase the effectiveness of ECGI, we provide new ways to visualize associated measurement and modeling errors. In this paper, we study source localization uncertainty in two steps: First, we perform Monte Carlo simulations of a simple inverse ECGI source localization model with error sampling to understand the variations in ECGI solutions. Second, we present multiple visualization techniques, including confidence maps, level-sets, and topology-based visualizations, to better understand uncertainty in source localization. Our approach offers a new way to study uncertainty in the ECGI pipeline.

## 1. Introduction

To rapidly diagnose heart disease, clinicians rely on the electrocardiogram (ECG), which records voltages on the torso surface. The voltages vary in response to changes in the heart’s electrical activity. Although the ECG quickly provides clinicians with information on abnormal rhythms or *arrhythmias*, it cannot reveal localized high-resolution spatial information about the heart’s electrical impulses.

For example, in arrhythmias involving added abnormal beats, such as premature ventricular contraction (PVC), a region of cardiac tissue initiates pathological heartbeats, thereby increasing a patient’s risk of sudden death (Messineo 1989). The lack of high-resolution spatial information from the ECG in locating this region is problematic, because one method of therapy involves a clinician locating and destroying the region through an invasive interventional procedure called catheter ablation. A catheter ablation procedure may last several hours with a frequently high rate of recurrence of the arrhythmia (Arya et al. 2010; O’Donnell et al. 2003).

Electrocardiographic imaging (ECGI) is one promising technique for increasing the speed and accuracy of ablation therapy. ECGI combines a patient’s computed tomography (CT) and magnetic resonance imaging (MRI) images along with the ECG to create a functional imaging modality (Johnson 1997; MacLeod et al. 2009; van der Graaf et al. 2014; Ghosh et al. 2008b). Challenges in ECGI may be categorized as technical (e.g., regularization, filtering techniques, and postprocessing methods), pathological (i.e., ability to extract features applicable to a specific pathology or arrhythmia), and clinical (i.e., benefits with respect to daily clinical practice) (Cluitmans et al. 2018). Our work addresses both the technical and clinical aspects of ECGI, with particular emphasis placed on using visualization techniques to better understand ECGI simulation uncertainty and aid in clinical decision-making.

Whereas the ECG may be thought of as a forward problem that relates the heart’s electrical activity to the recorded torso surface voltages, potential-based ECGI is the corresponding inverse problem that relates ECG measurements to heart surface voltages (Johnson 1997; Wang et al. 2011a; Rudy 2013). The mapping between the heart surface voltages and torso surface voltages may be written mathematically as

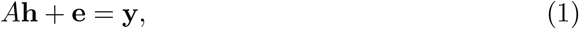

where *A* is a transfer matrix relating the heart surface voltages **h** to the torso surface ECG recordings **y**. The noise term **e** is modeled as a Gaussian distributed random variable to characterize uncertainties arising from multiple factors, e.g., model inaccuracies and sensor errors. The addition of such random error provides a more realistic representation of ECG measurements **y**, and hence, a more realistic representation of inverse solutions. In Equation (1), we added Gaussian noise **e** as a percent *p* of the ground-truth ECG torso surface observations, **y**\*, as

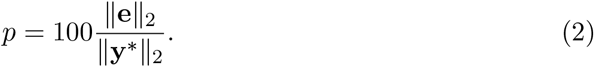

The forward problem estimates the torso surface potential **y** given **h**, and the inverse problem estimates **h** given **y**.

In recent years, researchers and clinicians have used ECGI to study a variety of arrhythmias, including reentrant pathways (Ghosh et al. 2008a) and ectopic heart beats (Wang et al. 2011c). ECGI may improve ablation therapy, but researchers do not have a good understanding of how small errors arising from ECG measurements, geometric approximations from imaging, and modeling assumptions for solving the underlying equations affect source localization in ECGI. Recent work by Tate et al. (2021) on quantifying geometric uncertainty resulting from variations in segmentation has shown some correlation between pericardial potential reconstructions and segmentation variability except in the posterior region of the heart.

Understanding uncertainties relevant to computational pipelines is a top research challenge in medical visualization (Ristovski et al. 2014; Karayiannis et al. 2004; Athawale et al. 2019; Fikal, Najib et al. 2019), as well as the visualization research field in general (Johnson and Sanderson 2003; Brodlie et al. 2012). Recently, visualizations were proposed by Burton et al. (2013) to study uncertainty associated with cardiac forward and inverse problems for 3D volumetric data. In our work, we explore new techniques to visually analyze and understand the uncertainty in epicardial surface data from ECGI simulations.

Building on preliminary work for source localization by France and Johnson (2016), we apply iterative Krylov methods with Monte Carlo error propagation to study the impact of ECG measurement errors on inverse solutions. We then propose a framework for deriving source localization confidence interval (CI) regions, and present applications of level-set and topology-based visualizations to visually analyze uncertainty in source localization. We propose that our CI visualizations could provide a sequential search strategy for clinicians in locating pathological heart beats during ablation therapy. Our level-set and topology-based visualizations can be useful in performing qualitative assessment of inverse solutions and extracting likely source positions.

We organize our paper as follows: In Section 2, we outline the mathematical framework for formulating and solving the inverse problem of electrocardiography. Section 3 discusses our Monte Carlo propagation strategy for studying the uncertainty of ECGI solutions. Section 4 describes our algorithms for developing probability maps and CI regions, as well as applications of level-set and topology-based techniques, for studying uncertainty in source localization. In Section 5, we show our results and discuss their implications. Finally, in Section 6, we present a summary and propose future work.

## 2. Inverse Problem of Electrocardiography

Here we describe the mathematical model, challenges in solving the inverse problem, our regularization algorithms, and the simulation setup.

### 2.1. Mathematical Model

The potential-based forward and inverse problems of electrocardiography are typically modeled using Laplace’s equation (Johnson 1997, 2015; Wang et al. 2011a,b). In our mathematical model, the potential *u* within a torso is modeled as a function of position **x** as

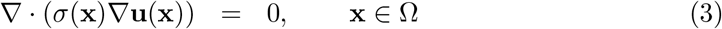

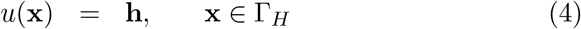

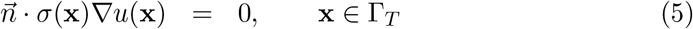

where Ω refers to the torso volume, Γ_*H*_ denotes the epicardial surface, and Γ_*T*_ indicates the torso surface. In this formulation, σ(**x**) is the electrical conductivity tensor, and 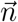 refers to the unit normal pointing outward from the torso surface, with Equation (5) stating no electric flux leaves the body into the air (Johnson 1997; Wang et al. 2011a, 2011b). Since our main contribution is visualizing uncertainty, we have implemented a simplified inverse model. We note that the visualization techniques we illustrate can be applied to any ECGI pipeline.

Several numerical methods exist for solving Equations (3–5). In this study, we used the finite element method to solve Equations (3–5) and rearranged the resulting stiffness matrix to form the transfer matrix *A* as described previously in Wang et al. (2011a,b) and Johnson (1997, 2015) to generate Equation (1).

### 2.2. Ill-Posedness and Ill-Conditioning

In our study, Equation (1) suffers from the ill-posedness common to inverse problems. Equation (1) is ill-posed because small changes in the observed ECG torso surface recordings lead to correspondingly large changes in the reconstructed heart surface potentials. In the discrete approximation, matrix *A* is highly ill-conditioned, and the singular values of *A* decay rapidly toward machine precision. Consequently, performing inversions using conventional routines greatly amplifies the impact of any numerical or measurement errors (Wang et al. 2011a, 2011b). To overcome the challenges associated with the ill-posedness and ill-conditioning, researchers employ regularization (Hansen 2010; Borràs and Chamorro-Servent 2021). In this paper, to increase the speed and scalability for Monte Carlo sampling, we used iterative methods to regularize solutions in Equation (1), as we describe next.

### 2.3. Regularization

In this study, we applied iterative regularization using the conjugate gradient least squares (CGLS) and preconditioned CGLS (PCGLS) methods. Milanič et al. (2014) have shown that the CGLS method performs as well as the standard Tikhonov methods in solving the ECG inverse problem with single dipole sources, but with the advantage of being computationally more efficient.

#### 2.3.1. Conjugate-Gradient Least Squares (CGLS)

The conjugate gradient least squares (CGLS) algorithm seeks the regularized solution after *k* iterations, **h_k_**, as demonstrated by Hestenes and Stiefel (1952) and Hansen (2010):

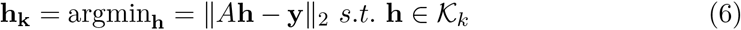

where 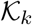 represents the *k^th^* Krylov subspace, which is formally defined as

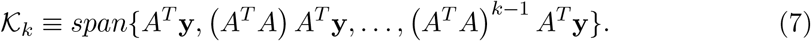

Starting from the zero vector at iteration zero, **h_0_**, this algorithm applies one multiplication with *A* and *A^T^* per iteration. The solution is formed as a linear combination of the Krylov vectors, and it becomes increasingly enriched in the direction of the principal eigenvector of *A^T^A* (Hansen 2010).

#### 2.3.2. Preconditioned Conjugate-Gradient Least Squares (PCGLS)

We also solved Equation (1) with the preconditioned CGLS (PCGLS) algorithm, using the Laplacian operator *L* over the heart surface as the right preconditioner. The matrix *L* was formed as described in Huiskamp and van Oosterom (1988). Because *L* has a nontrivial null-space *W*, the PCGLS method requires formation of the A-weighted pseudo-inverse of *L* (Hansen 2010),

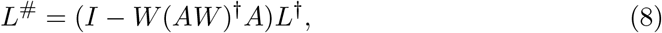

where *I* is the identity matrix. The component of **h_k_** that exists in the null space of L is given as

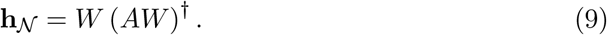

Then, defining 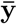 as

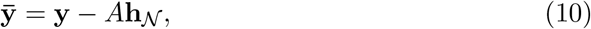

and with *Ā* = *AL*#, we solved 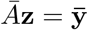 for **z_k_** using the traditional CGLS routine. The PCGLS solution can then be obtained as

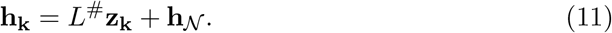

As in the CGLS routine, the PCGLS algorithm terminates at some iteration to prevent under-regularization (Hansen 2010).

#### 2.3.3. Choosing the Iteration Parameter k

In choosing the solution at which to stop iterations, we used both the norm of the residual and the norm of the solution. In using the norm of the residual, we used the Morosov discrepancy principle, in which we stopped the iterations as soon as the norm of the residual was approximately equal to some constant *γ* times the norm of the noise ║**e**║_2_, or *γ*║**e**║_2_ (Hansen 2010; Kaipio and Somersalo 2004). Following the example of the study by Calvetti et al. (2015), we used *γ* = 1.2.

However, as we discuss in our results in Section 5, the discrepancy principle may severely under-regularize the solution when the modeling error is significant relative to the measurement error. To address this under-regularization, we used physiologically based mathematical constraints for the norm of the solution in limiting the termination iteration *k* for the CGLS and PCGLS algorithms. Specifically, previous studies on cardiac electrograms recorded and derived relationships on the scalar gain *G*, between the *ℓ*_2_ norm of the torso and heart voltages at a moment in time, or ║**h**║_2_ *G*║**y**║_2_.

These studies had an equal number of torso and heart nodes (i.e., *m* = *n*) (Davenport et al. 1995; Davenport 1995). To account for differences in the number of heart and torso nodes in this study, we used a slightly modified formula of 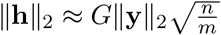 and a gain value **G** of 7.6, a value slightly less than the maximum experimentally derived value from the *ℓ*_2_ norm in the scalar gain studies (Davenport et al. 1995; Davenport 1995). Putting restrictions on the norm of the solution prevents under-regularization, particularly when the modeling error exceeds external noise error. Furthermore, slight over-regularization in inverse reconstructions seems to be preferred for clinical applications as opposed to any form of under-regularization (Milanič et al. 2014).

### 2.4. Simulation Setup

Figure 1 illustrates the heart in torso geometry (left) and the 250-uniform-ECG-electrode-measurement configuration (right) used in this study. Other studies use a similar electrode configuration and number of electrodes (Ghosh et al. 2008a; Rudy 2013).

**Figure 1.**
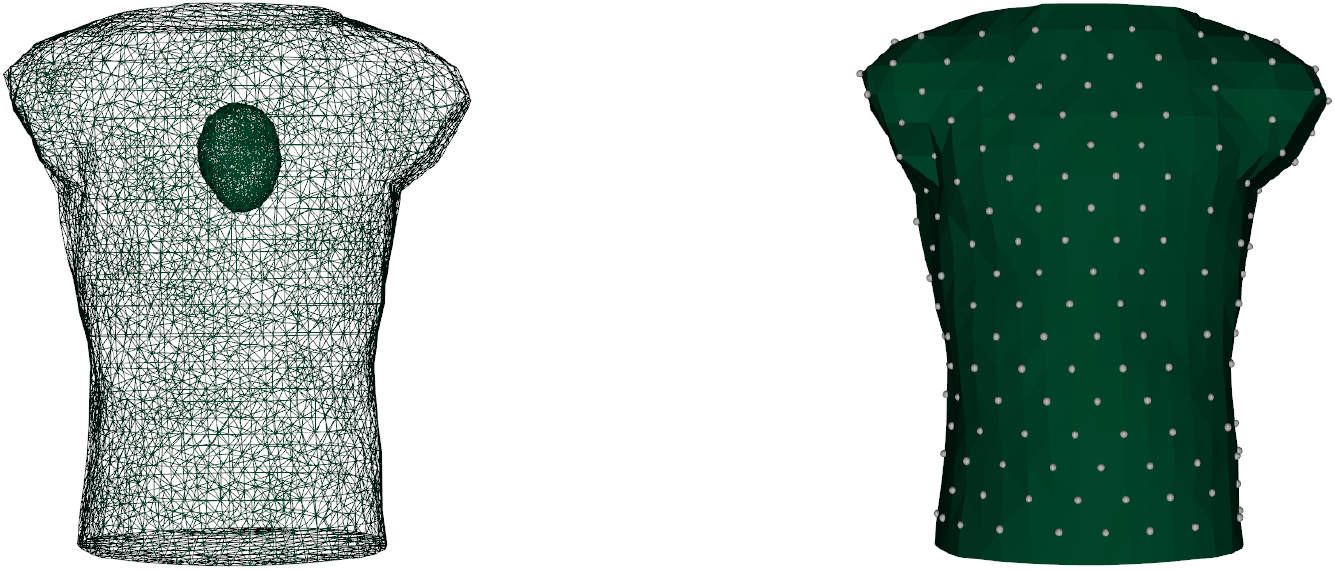
Inversions utilized the heart torso geometry (left) with a 250-uniform-lead configuration (right).

For this study, we added noise from 0.01% to 3%, values similar to those found in other studies (Wang et al. 2011a, b, 2013; Burnes et al. 2000). Additionally, we used a different transfer matrix *A* in forming the ground-truth observations **y**^*^ compared with the transfer matrix used in inversions to avoid so-called “inverse crime”, where the solution is biased by using the same mesh for forward and inverse simulations (Kaipio and Somersalo 2007). For the forward model simulation, we utilized a higher resolution finite element model as illustrated in Table 1. We use a single stimulation point for our analysis throughout the paper. Additionally, for our inverse simulations, we added 2 mm Gaussian geometric error to the torso surface recording sites, as in other studies (Burnes et al. 2000).

**Table 1.** Forward and Inverse Model Resolution

| Number of Attributes | Forward Model | Inverse Model |
| --- | --- | --- |
| elements | 128,898 | 75,605 |
| heart nodes | 3,573 | 2,206 |
| torso nodes | 7,328 | 4,754 |

## 3. Monte Carlo Approach to Studying Solution Uncertainty

Having obtained an initial solution **h_k_** using the CGLS or PCGLS routine, we forward-propagated the solution to form an assumed noise-free right-hand side 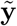 with

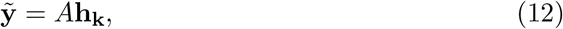

as described in Aster et al. (2013). To perform Monte Carlo error analysis, we sampled a noisy solution 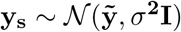, where *σ* represents the standard deviation of the noise, a value that was fixed to produce errors at the same percentage *p* as in the original Equation (1). We then used the CGLS and PCGLS routines to obtain individual inversion samples, just as we obtained **h_k_** in originally solving Equation (1) (Aster et al. 2013; France and Johnson 2016). We obtained an ensemble of 200 Monte Carlo samples per simulation.

## 4. Visualizing Source Localization Uncertainty

We analyzed the uncertainty in source localization across 200 Monte Carlo samples per simulation via probability maps, confidence interval regions, level-set visualizations, and topology-based visualizations.

### 4.1. Probability Maps

For probabilistic maps, similar to the early study by France and Johnson (2016), we located the top 3% of the lowest voltage values (with the lowest voltage denoting the source (Wang and Rudy 2006)), and averaged these locations over the 200 samples to form a probabilistic representation for source localization. In France and Johnson (2016), probabilistic maps were visualized with direct mapping of probability to opacity. In our probabilistic map visualizations, we used color maps to segment the regions of high probability (> 0.75), moderate probability (between 0.5 and 0.75/ between 0.25 and 0.5), and low probability (< 0.25). We propose that the segmented visualizations can potentially benefit clinicians in performing sequential searches for source localization.

### 4.2. Confidence Intervals

Although probability maps represent the probability mass function for source localization, the confidence interval regions may be considered as the corresponding cumulative density function for source localization. To generate these visualizations for confidence intervals, we performed integration of the probability maps from the position of the estimated source location. First, we determined the estimated source location, along with the probability maps. For each node, we assigned the confidence interval value at a particular node *i*, which is the value of the sum of the probabilities that reside within the radius from the estimated source location to that node *i*. After calculating the confidence interval values at each node, we divided the 25%, 50%, and 75% confidence interval regions using contour lines via the marching triangles algorithm (Hilton and Illingworth 1997). Again, the bands visualized with confidence interval visualizations can potentially help clinicians perform sequential searches for source localization.

### 4.3. Level-Set Visualizations

We perform level-set visualizations to identify the regions that could contain the source of arrhythmia. Level-sets (Lorensen and Cline 1987) are a fundamental surface-based visualization technique for gaining insight from complex scientific data. Mathematically, for a function 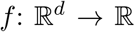 defined on a d-dimensional manifold, its level-set *S* for isovalue *c* is defined as 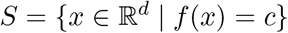. Figure 2b illustrates level-sets of a synthetic Ackley function (Ackley 1987) shown in Figure 2a. The level-sets in Figure 2b for different isovalues *c* are displayed in different colors. In the ECGI context, function *f* is a mapping from nodes of a mesh representing the heart surface to inverse solutions denoting epicardial potentials. We investigate level-sets with relatively low isovalues to understand potential positions of arrhythmia.

**Figure 2.**
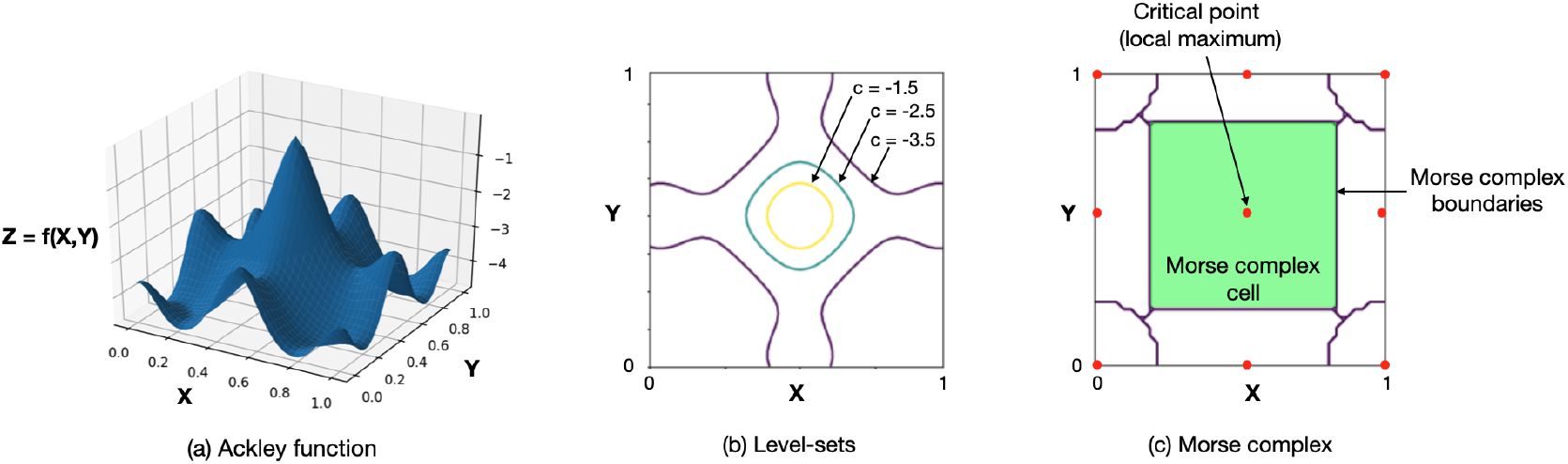
Illustration of level-sets in image (b) and Morse complexes in image (c) for the synthetic Ackley function *f* depicted in image (a). Level-sets are visualized for different isovalues *c*. The Morse complex visualization denotes nine cells (a single cell highlighted in green) corresponding to nine critical points (local maxima indicated by red dots) of the Ackley function. Gradients within a single cell flow to its corresponding critical point.

We guide our selection of an isovalue for level-set visualizations using parallel coordinate (Inselberg 1985) and histogram plots. Parallel coordinate plots are a well-known visualization technique to study correlation among dimensions of multivariate data. For our analysis, in the parallel coordinate plot, we treat each node of a mesh representing the heart surface as a single dimension of a parallel coordinate plot. We then plot the potentials across all nodes and 200 samples. Likewise, in the histogram plot, cardiac potentials across all nodes and 200 samples are grouped into bins. We then use these plots to gain insight into the relatively low potential values that are observed across all samples. We pick one of the low potential values as the isovalue for level-set rendering. Level-sets with relatively low isovalues help us extract regions that correspond to a potential source (Wang and Rudy 2006). Finally, we visualize isocontours for the selected isovalue using spaghetti plots (Potter et al. 2009) and isocontour variation plots (Whitaker et al. 2013).

### 4.4. Topology-Based Visualizations

Topological data analysis is a powerful tool for understanding complex simulation datasets (Miller et al. 2006; Bremer et al. 2010). We propose visualizations of topological abstractions, specifically, critical points and Morse complexes (Edelsbrunner et al. 2001), of CGLS and PCGLS inverse solutions to gain insight into the likely source positions and their variations. Let function 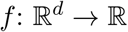 be defined on a d-dimensional manifold, and let ∇*f* denote its gradient field. A point *x* on a manifold is considered critical if ∇*f* = 0. Given a Morse function *f* defined on a d-dimensional manifold, i.e, a function with no flat regions, the Morse complex of *f* decomposes the manifold into regions (referred to as cells) with uniform gradient behavior. Figure 2c illustrates the Morse complex segmentation of the Ackley function shown in Figure 2a. In Figure 2c, nine Morse complex cells correspond to nine critical points of (local maxima) of the Ackley function. In our case, the Morse complexes segment the heart surface into cells, where gradients within a single cell terminate in a single local minimum associated with a cell (also known as an ascending manifold). Thus, local minima of ECGI solutions provide insight into the positions that have the smallest potential within their local neighborhood (represented by the Morse complex cell), thus indicating potential source positions.

## 5. Results and Discussions

We now present results for each visualization technique described in Section 4 to understand the source localization uncertainty.

### 5.1. Probability and Confidence Maps

We first present the uncertainty visualization results using probability maps and our proposed confidence maps in Figure 3. Figure 3a visualizes the ground-truth heart surface voltages (in mV). In all plots, the white dot indicates the position of the ground-truth source. Figures 3b-c visualize the CGLS and PCGLS inverse solutions at 1.5% external noise for a single Monte Carlo sample.

**Figure 3.**
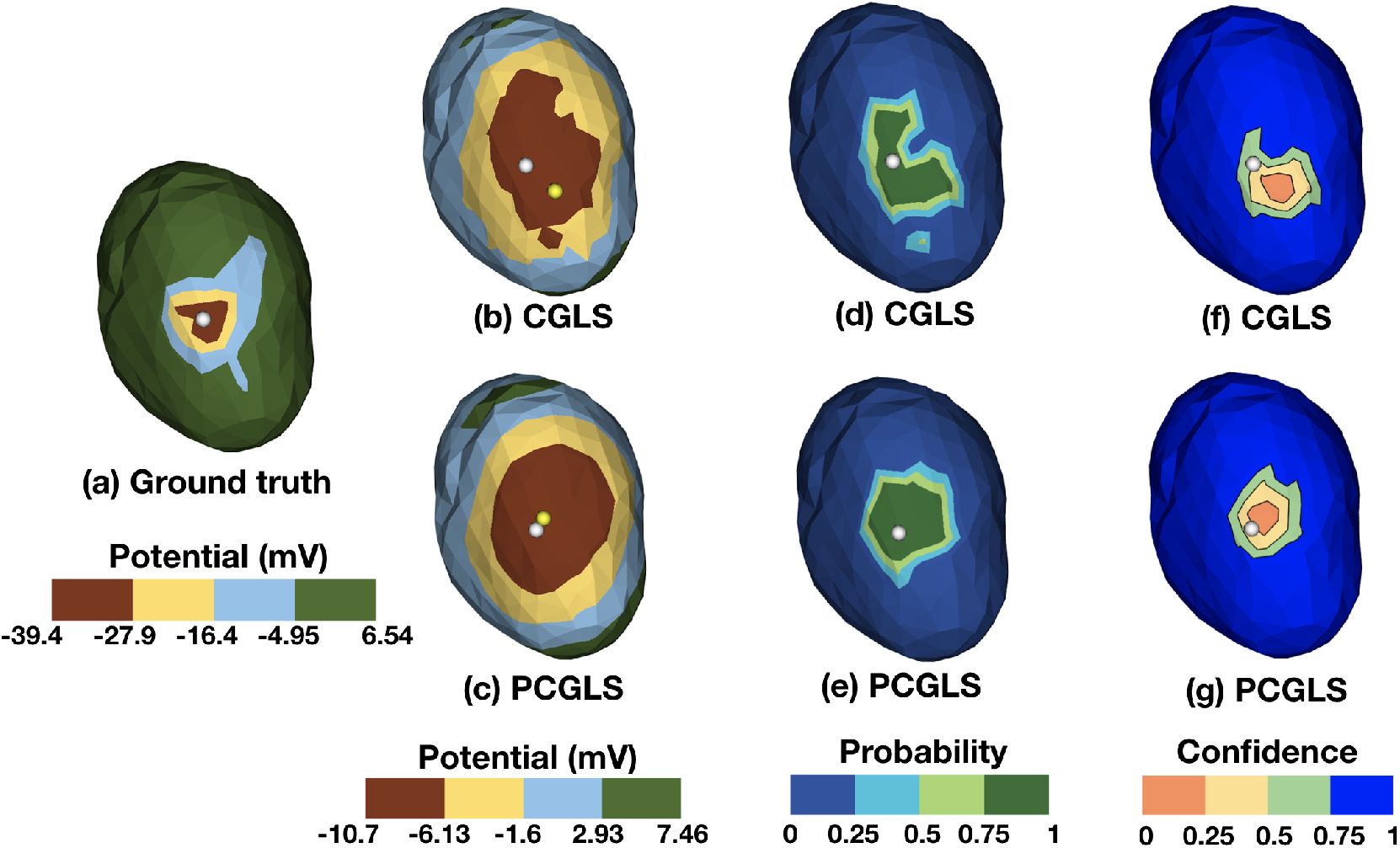
Uncertainty analysis of Monte Carlo simulations with probabilistic and confidence maps: (a) The ground-truth voltages, (b,c) the CGLS and PCGLS inversions for a single Monte Carlo sample at 1.5% external noise, (d,e) probabilistic maps, (f,g) confidence maps. The white dots denote the true source positions, whereas the yellow dots in (b,c) denote the estimated source positions. The results (a-c) are colormapped with the potential values, the results (d,e) are colormapped with the probabilities, and the results (f,g) are colormapped with the confidence values.

Figures 3b-c illustrate the challenges in obtaining inverse solutions in ECGI, i.e., the reconstructed voltages in Figures 3b-c differ significantly in both range and magnitude in comparison to the ground-truth, Figure 3a. This difference is evidence of the significant ill-conditioned nature of discrete ECGI problems, as discussed in Section 2.2. Inversions in Figures 3b-c were performed with the lower resolution “Inverse Model” mesh in Table 1, and with additive 2 mm Gaussian geometric error on the heart surface, as described in Section 2. The yellow dot in Figures 3b-c indicates the estimated source location for each individual visualization. This yellow dot corresponds to the global minima of the reconstructed epicardial potentials.

Figures 3d-e and 3f-g illustrate the probability maps (France and Johnson 2016) and confidence maps, respectively. For the probability maps visualized in Figures 3d-e, the darker green areas indicate the regions of higher probability for source localization. In the confidence maps visualized in 3f-g, the contour lines separate the 25% (orange), 50% (yellow), and 75% (light green) CI regions for source localization. Specifically, in the orange regions, our model states that the probability of finding the source is less than or equal to 25%. Likewise, the probability of finding the source in the combined orange and yellow regions is less than or equal to 50%. Note the lack of symmetry around the sites of stimulation, which is due to the coarseness of the grid and its unstructured nature.

The green region representing the high source localization probability in probabilistic maps (Figures 3d-e) still covers a relatively large surface area. The CI visualizations could provide clinicians with an improved sequential search strategy for planning ablation therapy. For example, during an ablation procedure, a clinician might sequentially search the 25%, 50%, and 75% CI regions to locate the source of cardiac tissue responsible for spontaneous pathological heart beats or reentrant wave activity.

The vertex positions of confidence/probability map contours can be utilized to quantify the positional uncertainty of source localization. For example, in Figure 3g, the maximum Euclidean distance between contour vertices for the 25% confidence interval is 8.56 mm. The error in source localization, thus, may not exceed 8.56 mm, assuming that the 25% confidence region is locally planar and the source resides in the 25% confidence region. More advanced techniques of error quantification may be developed in the future that take into account contour vertex positions and heart surface curvature.

Figure 4 illustrates probability maps for the CGLS and PCGLS routines at various external noise levels, as defined in Equation (2). In both algorithms, the probability tended to aggregate around the ground-truth source location until approximately 3% external noise. Figure 5 illustrates the corresponding CI regions for the CGLS and PCGLS routines. For the CGLS and PCGLS routines, the ground-truth source location fell within the 50% CI region (yellow) and 25% CI region (orange), respectively, for all noise levels.

**Figure 4.**
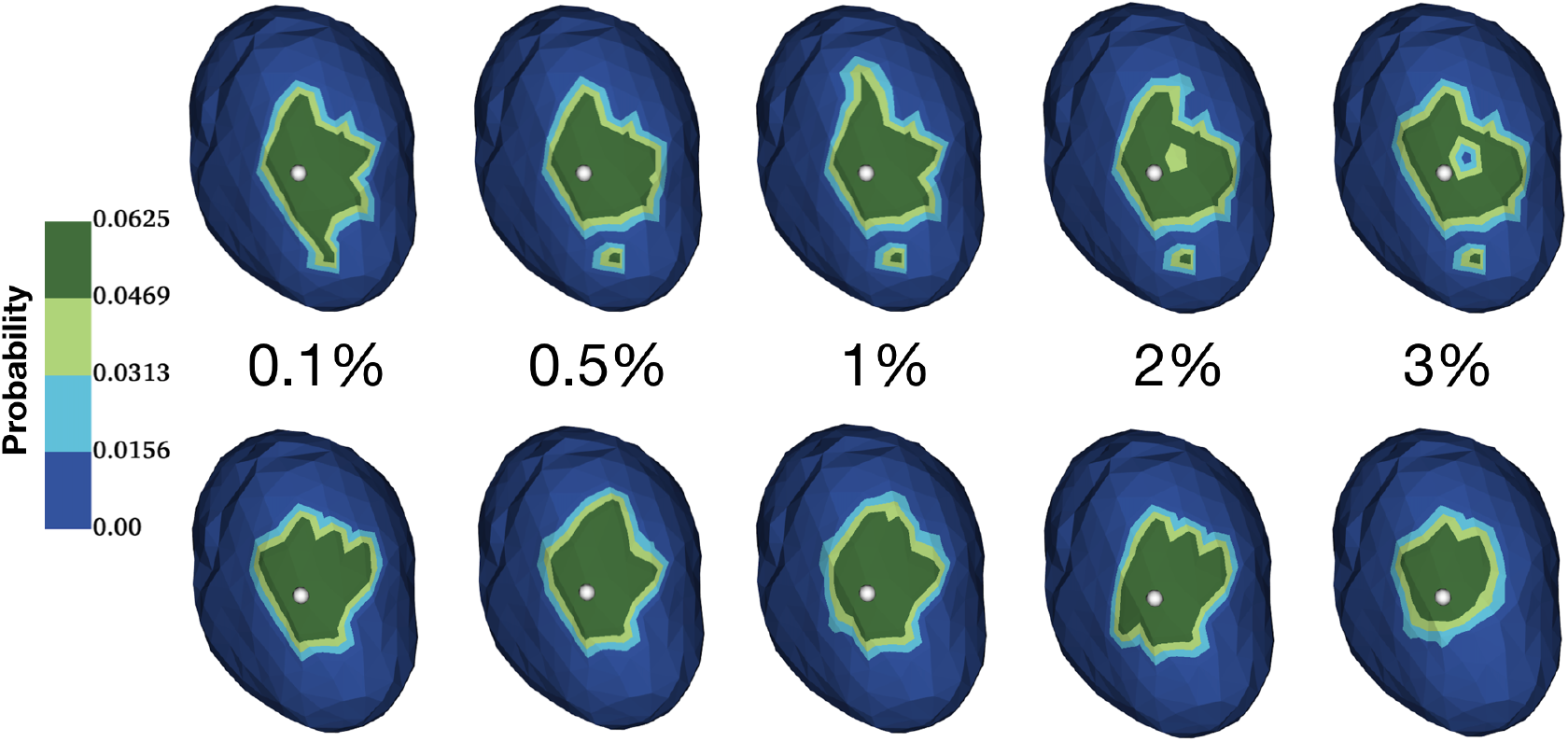
Probability maps for source localization illustrate uncertainty as a function of noise in ECG observations for the CGLS inversion (top row) and PCGLS inversion using a Laplacian preconditioner (bottom row). White dots mark the ground-truth source location.

**Figure 5.**
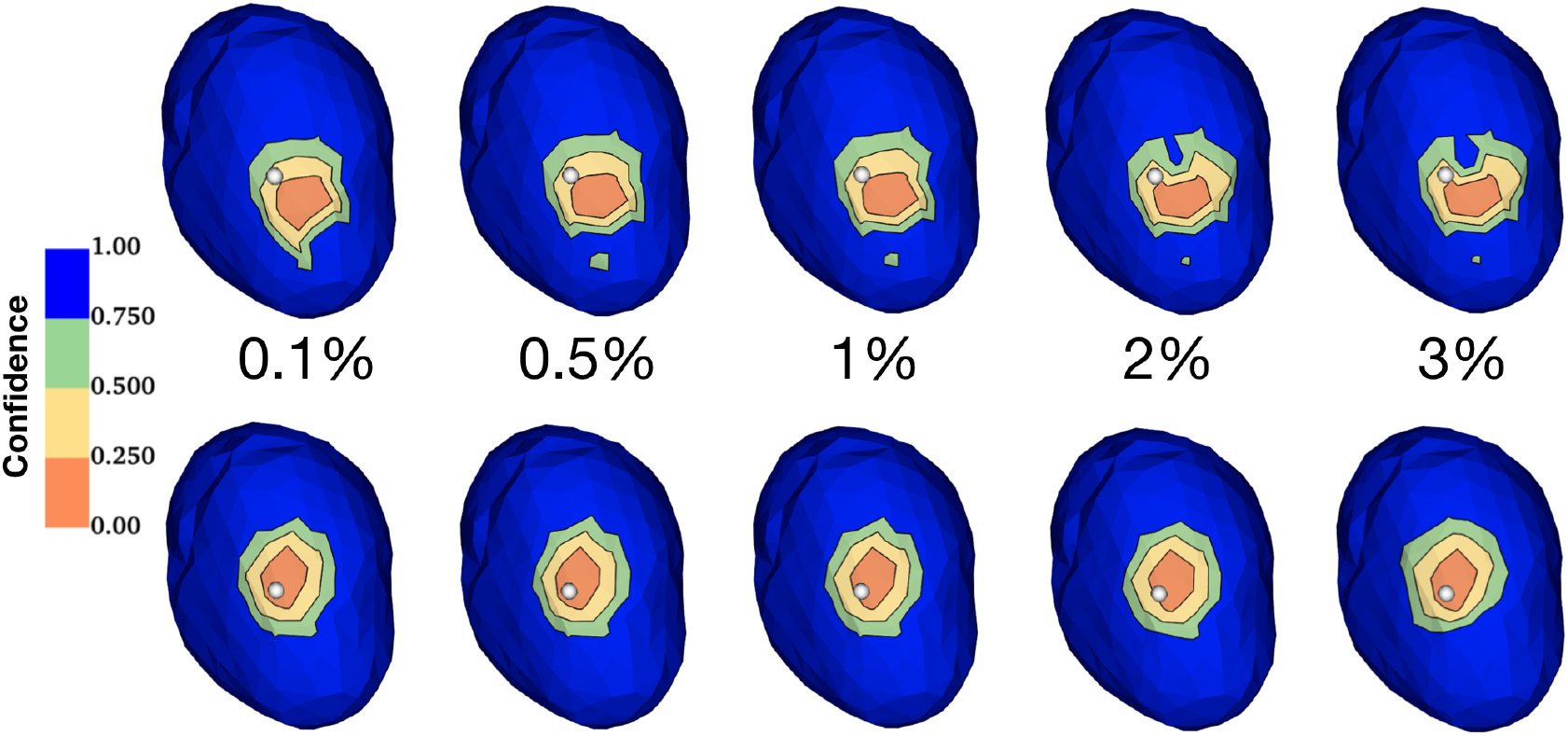
Confidence interval (CI) regions for source localization illustrate uncertainty as a function of external noise in ECG observations for the CGLS inversion (top row) and PCGLS inversion using a Laplacian preconditioner (bottom row). Dots mark the ground-truth source location. Contour lines separate the 25%, 50%, and 75% CI regions.

Figure 6 illustrates the convergence of the CGLS routine at 0.5% and 2.5% external noise levels, both with and without modeling error using the CGLS algorithm. Although not shown, PCGLS gives similar results. At 0.5% external noise (Figure 6, left), the norm of the residual approaches the norm of the noise after 20 iterations to satisfy the Morosov discrepancy principle. However, in the presence of modeling error in the transfer matrix *A* (from a coarser mesh resolution and torso surface electrode position errors, as described in Section 2), the CGLS residual does not converge to the norm of the external noise, even after several hundred iterations (not shown). At 2.5% noise, however (Figure 6, right), the convergence patterns of CGLS on transfer matrices with and without modeling error follow a nearly identical pattern. Figure 6 (left) illustrates the need to also use the norm of the solution as a constraint to prevent under-regularization, especially when the ratio of modeling error to external error is large. Other studies also report difficulties in regularization when modeling error exceeds measurement error (Johnston and Gulrajani 2002).

**Figure 6.**
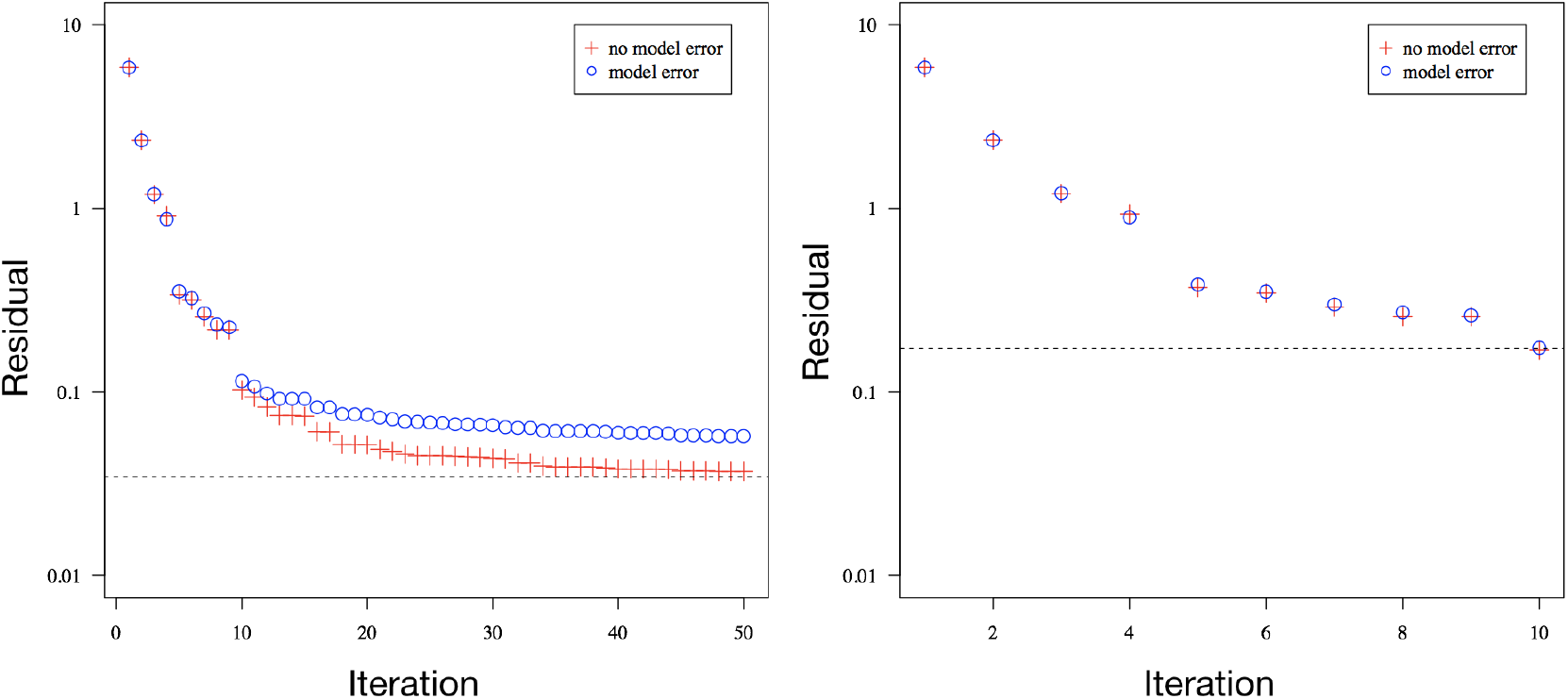
At 0.5% external noise (left), modeling error affects convergence, whereas at 2.5% external noise (right), convergence appears similar both with and without modeling error. For both plots, the dashed line indicates the Morosov discrepancy principle termination criterion.

### 5.2. Level-Set Visualizations

Next, we study the source localization uncertainty by extracting level-sets of epicardial potentials, in which we guide the isovalue selection with parallel coordinate plots and histograms. Figure 7 shows a parallel coordinates plot (left) and a histogram (right) plot of the 200 samples denoting the epicardial voltage recordings at each of the 337 heart nodes represented at the 0.5% noise level. A parallel coordinate plot has the advantage of not losing spatial context as opposed to a histogram. These plots were useful in the visualization and selection of the low-magnitude heart surface voltages corresponding to the sources, as described in Section 4. We used these plots in choosing the isovalue −6.5mV.

**Figure 7.**
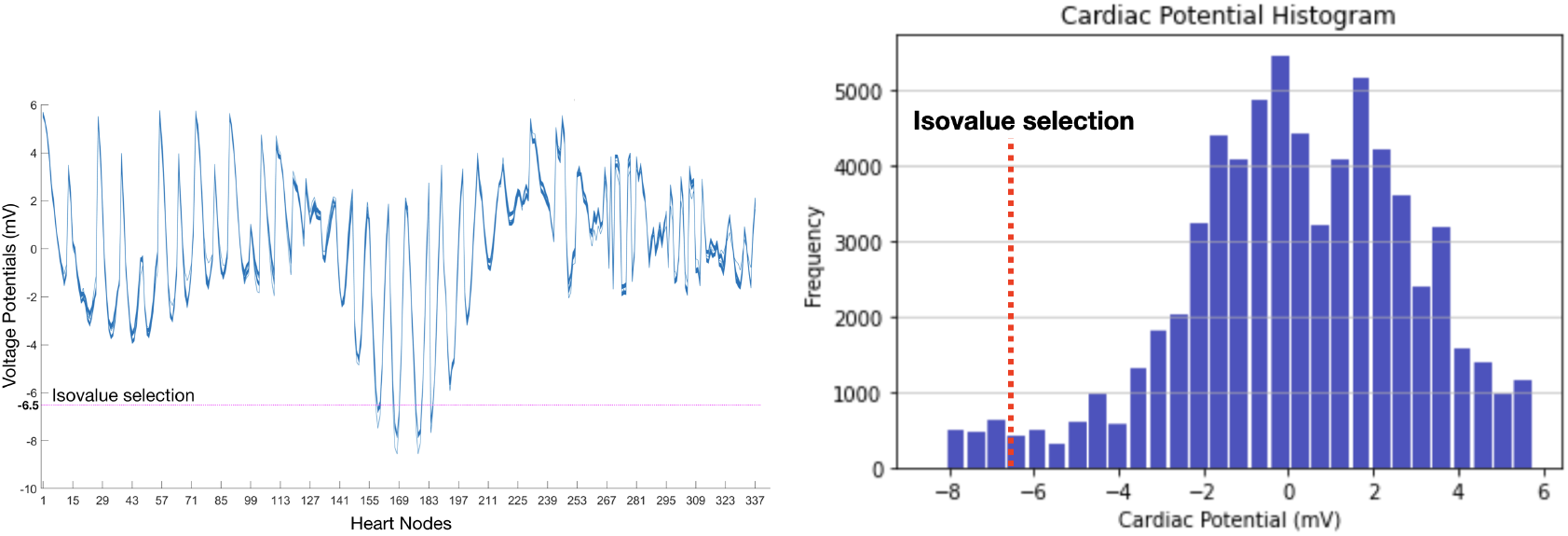
Parallel coordinate (left) and histogram (right) plots of heart surface voltage recordings from 200 Monte Carlo samples at the 0.5% noise level. The dotted lines indicate our isovalue selection (−6.5mV) for level-set rendering.

Figure 8 shows a spaghetti plot of level-set renderings for the isovalue −6.5 mV across different noise levels. As can be observed, the ground-truth source location fell within the confines of level-sets for all noise levels. Thus, the region represented by the outermost level-set of a spaghetti plot denotes the likely existence of a source location. In Figure 8, the isocontour shape for the PCGLS solutions (bottom row) appears more closely aligned with the ground-truth isocontour (purple) shape when compared to the shape of the CGLS isocontours (top row).

**Figure 8.**
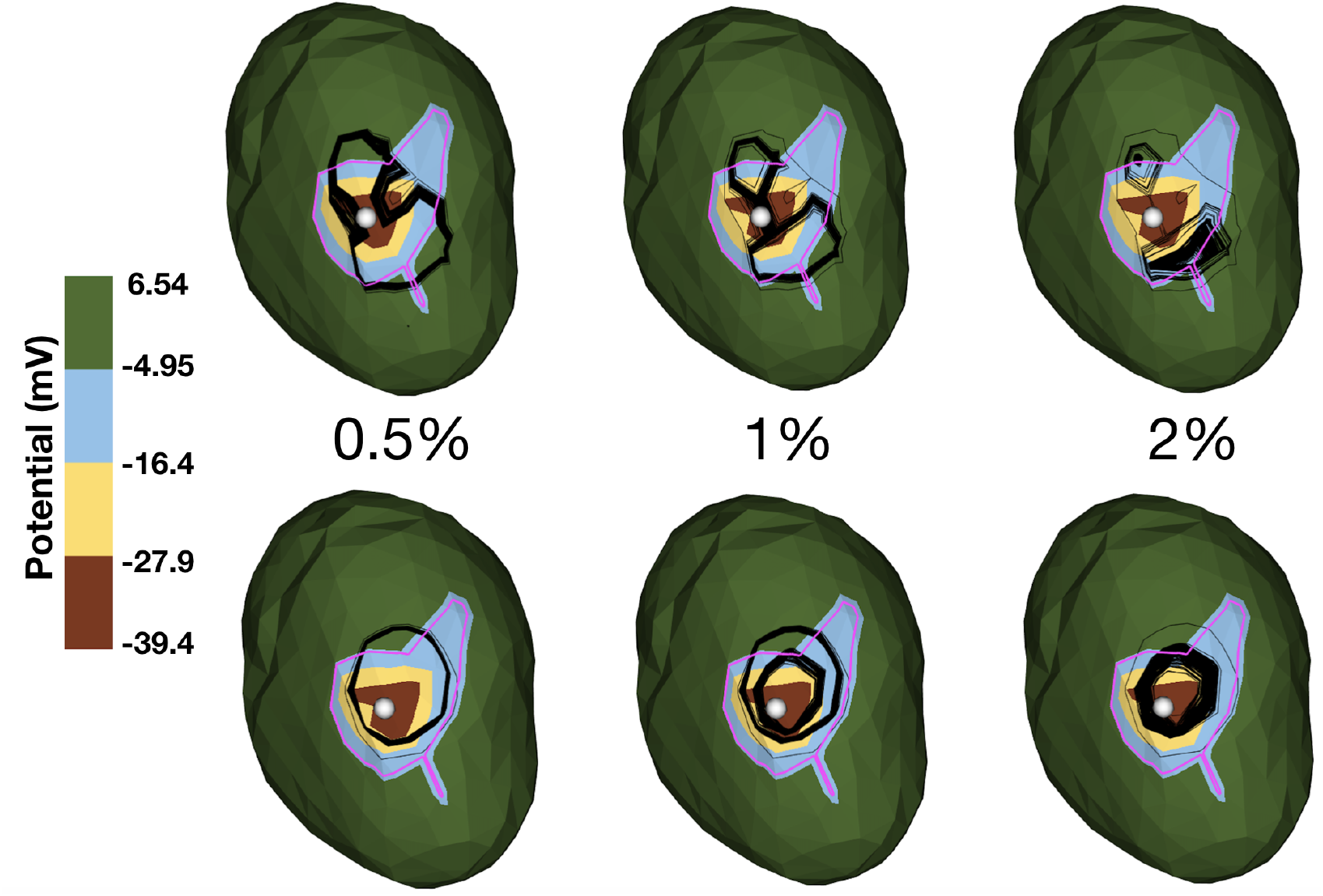
Isocontour maps for sample *isovalue* = −6.5 mV showing the ground-truth isocontours (purple) overlaid with isocontours (black) from the 200 samples. The top row shows isocontours without any smoothing (i.e., CGLS) whereas the bottom row shows isocontours with Laplacian smoothing applied (i.e., PCGLS). The columns indicate different noise levels. Dots mark the ground-truth source location.

We visualize an isocontour variation plot to reduce the clutter caused by spaghetti plots. Figure 9 visualizes the variations corresponding to the spaghetti plots shown in Figure 8. Figure 9 may be easier to analyze and provides a less cluttered visualization because the 200 samples are summarized by the minimum (white), maximum (red), and average (yellow) isocontours overlaid on the ground-truth (purple). The minimum isocontour refers to the isocontour generated by using the minimum value from the reconstructed voltage potentials across all simulations. Likewise, the maximum isocontour refers to the isocontour generated by using the maximum value from the reconstructed voltage potentials, and the average isocontour refers to the isocontour generated by using the average value from the reconstructed voltage potentials. These isocontours are then overlaid on the ground-truth. This approach significantly reduces clutter while clearly showing the variation, and therefore, uncertainty, contained in the reconstructed solutions.

**Figure 9.**
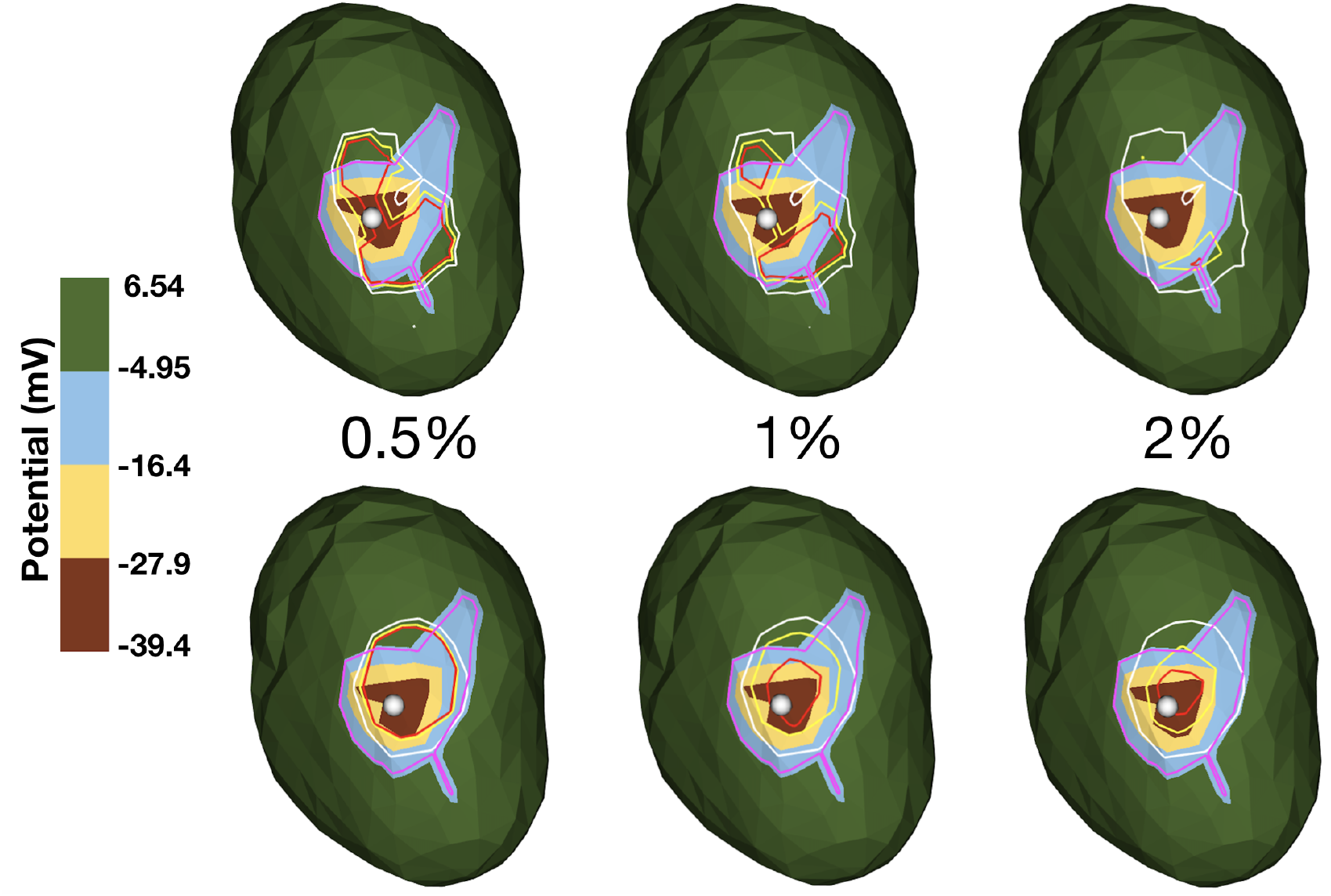
Isocontour maps for sample *isovalue* = −6.5 mV showing the ground-truth isocontours (purple) overlaid with the minimum (white), maximum (red), and average (yellow) isocontours from the 200 samples at each noise level. The top row shows isocontours without any smoothing (i.e., CGLS) whereas the bottom row shows isocontours with Laplacian smoothing applied (i.e., PCGLS). The columns indicate different noise levels.

### 5.3. Topology-Based Visualizations

Lastly, we analyze the source uncertainty via visualization of critical points and Morse complexes derived from ECGI solutions. Figure 10a visualizes the critical points (spheres) and Morse complex (white contours) for the ground-truth. Similar visualizations are produced for the CGLS (Figure 10b) and PCGLS (Figure 10c) solutions for the 1% noise level and a single sample. The visualizations are performed in ParaView (Ayachit 2015) using the topology toolkit (Tierny et al. 2018). The local minima of all three fields are visualized with the dark blue spheres, and the local maxima are visualized with the orange spheres. As indicated in Figure 9b, there are two local minima (enclosed by yellow dotted boxes), indicating uncertainty in source positions. However, this uncertainty can be further reduced by employing better regularization schemes, e.g., PCGLS, as shown in Figure 10c. Even though Figure 10 visualizes critical points for the sample 50, a similar trend is observed across all simulations.

**Figure 10.**
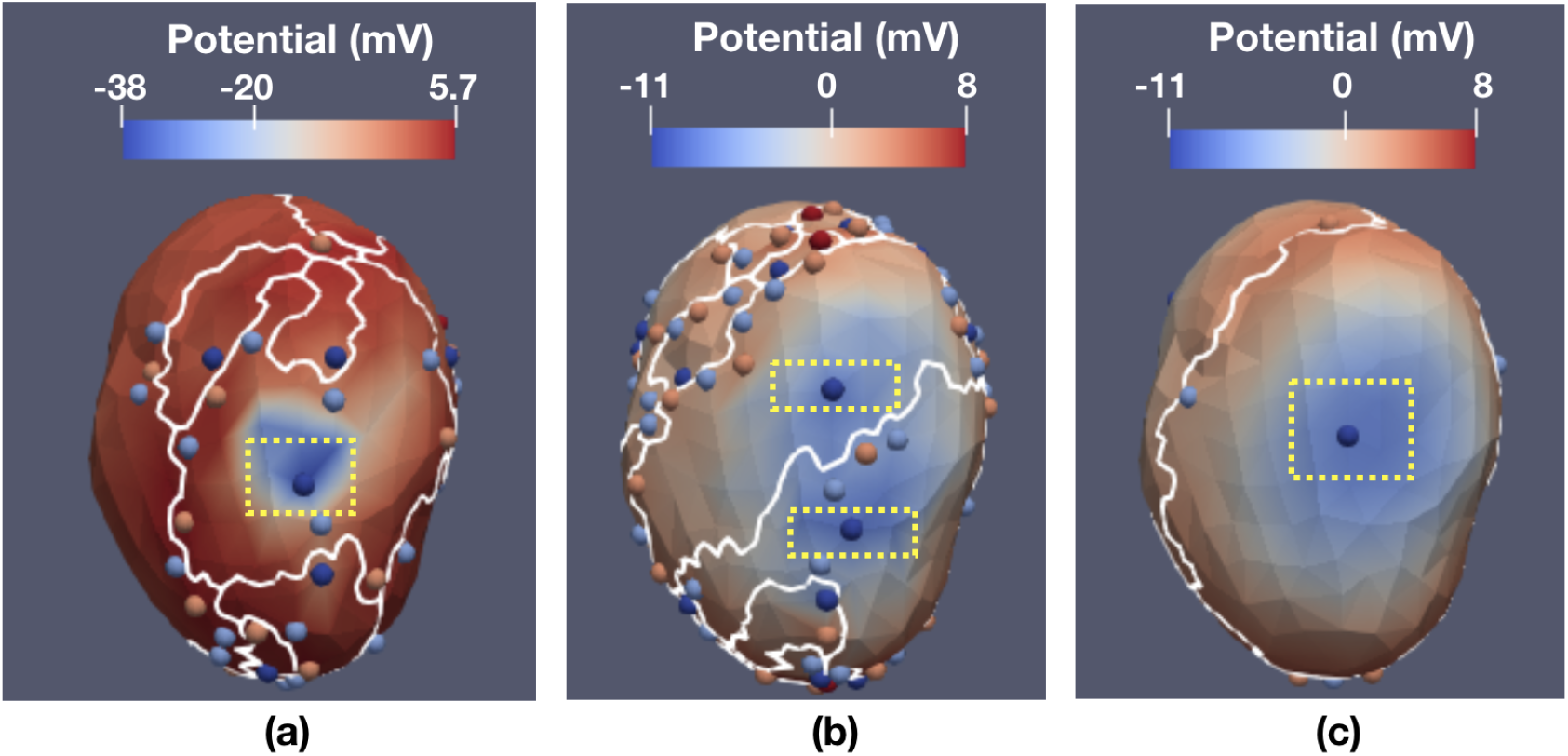
Visualization of Morse complexes and critical points for (a) ground-truth, (b) CGLS at 1% noise, and (c) PCGLS at 1% noise. The white contours denote the Morse complex cells, the dark blue spheres denote the local minima associated with each Morse complex cell, and the orange spheres denote local maxima of respective fields. The potential values are colormapped on the heart surface. The local minimum enclosed by the bottom yellow box in the CGLS visualization is situated closer (6.55 mm) to the true solution, indicated by the yellow box in the ground-truth, when compared to the PCGLS solution (7.72 mm).

## 6. Conclusion and Future Work

In this paper, we use multiple visualization techniques, several of which are new to ECGI applications, to study the impact of measurement and modeling errors associated with the ECGI pipeline on epicardial source localization. Specifically, we present applications of confidence maps, level-sets, and topology-based visualizations for effective analysis of uncertainty in source localization. In the future, we would like to study the sensitivity of source localization to variations in other ECGI parameters, such as electrical conductivity and number and configuration of ECG leads. We would also like to study and visualize uncertainties arising from multiple sources and add a quantitative metric to our probability maps and confidence maps to reflect the multiple localization errors resulting from multiple pacing site estimations. The analysis for multiple sources would require additional research into more sophisticated statistical models and uncertainty visualizations and would be an interesting extension of the methods presented here. Lastly, we would like to study how the different visualization methods we presented compare under different arrhythmia scenarios.

## Acknowledgments

This project was supported by grants from the National Institute of General Medical Sciences (P41 GM103545-18, R24 GM136986), the Intel Graphics and Visualization Institutes of XeLLENCE, and the Scientific Discovery through Advanced Computing (SciDAC) program in the U.S. Department of Energy. The authors would like to give special thanks to Professor Rob MacLeod for generously sharing the electrocardiogram data.

